# Integrating genomics and multivariate evolutionary quantitative genetics: A case study of multivariate constraints on sexual selection in *Drosophila serrata*

**DOI:** 10.1101/2021.08.09.455747

**Authors:** Adam J. Reddiex, Stephen F. Chenoweth

**Affiliations:** School of Biological Sciences, The University of Queensland, QLD 4072, Australia; Research School of Biology, Australian National University

**Keywords:** *multivariate quantitative genetics*, *multi-trait GWAS*, *sexual selection*, *genomics*, *Lande equation*

## Abstract

In evolutionary quantitative genetics, the genetic variance-covariance matrix, **G**, and the vector of directional selection gradients, ***β***, are key parameters for predicting multivariate selection responses and genetic constraints. Historically, investigations of **G** and ***β*** have not overlapped with those dissecting the genetic basis of quantitative traits. Thus, it remains unknown whether these parameters reflect pleiotropic effects at individual loci. Here, we integrate multivariate GWAS with **G** and ***β*** estimation in a well-studied system of multivariate constraint; sexual selection on male cuticular hydrocarbons (CHCs) in *Drosophila serrata*. In a panel of wild-derived resequenced lines, we augment genome-based REML, (GREML) to estimate **G** alongside multivariate SNP effects, detecting 532 significant associations from 1,652,276 SNPs. Constraint was evident, with ***β*** lying in a direction of **G** with low evolvability. Interestingly, minor frequency alleles typically increased male CHC-attractiveness suggesting opposing natural selection on ***β***. SNP effects were significantly misaligned with the major eigenvector of **G, *g***_max_, but well aligned to the second and third eigenvectors ***g***_2_ and ***g***_3_. We discuss potential factors leading to these varied results including multivariate stabilising selection and mutational bias. Our framework may be useful as researchers increasingly access genomic methods to study multivariate selection responses in wild populations.

## Introduction

There are two robust findings in evolutionary biology that are difficult to reconcile. The first, is the ubiquity of genetic variance in individual traits in nature evidenced by non-zero heritability estimates for almost all traits [1, 2] and ongoing responses to artificial selection [3]. The second is that traits are exposed to strong directional and stabilising selection [4-6]. The contradiction lies in the fact that traits under strong selection are expected to have low genetic variance [7], because when alleles are fixed or purged from the population, genetic variance is reduced. Furthermore, traits closely aligned to fitness tend to exhibit greater levels of genetic variance than metric traits [8]. How genetic variance is maintained in the face of natural selection is a fundamental question for theoretical population geneticists, however, the mutation-selection balance and balancing selection models proposed thus far have yet to convincingly reconcile the observed levels of genetic variance with known rates of mutation and strengths of natural selection found in nature [6].

Given high levels of genetic variance and the presence of strong directional selection, an intuitive prediction is that we would expect to observe constant evolution in natural populations. However, long-term studies of contemporary populations, do not support this suggesting that evolutionary stasis may be more common [9, 10]. This lack of evolutionary change suggests the presence of genetic constraints. Natural selection is unlikely to act on traits individually, but rather on multiple, often genetically correlated, traits simultaneously [11]. Geometric analyses of the additive genetic variance-covariance matrix, **G**, [12] have revealed uneven distributions of genetic variance across phenotypic space, with some trait combinations possessing little to no genetic variation [11, 13-15]. The observation that populations tend to diverge in directions of phenotypic space harbouring the greatest genetic variance is further evidence that for some directions in multi-trait space, there may be little to no genetic variance to facilitate evolutionary change [16, 17].

The basis for our understanding of how a population will respond to multivariate directional selection, defined by the vector of directional selection gradients, ***β*** [18], is derived from the Lande equation **Δ*Z* = G*β*** [12]. Lande’s equation elegantly reveals how the structure of **G** constrains evolution. For a vector of multivariate normal traits, ***Z, β*** represents the direction in trait space in which relative fitness has the steepest local rate of increase. Any deviation in **Δ*Z*** from ***β*** is a result of **G** rotating and scaling the evolutionary response sub-optimally [11]. The degree to which **G** constrains responses to selection can be characterised in multiple ways including the angle between **Δ*Z*** and ***β*** [19] or by measuring the degree to which **Δ*Z*** is reduced in length compared to ***β*** [20].

The utility of **G** to assess longer-term constraints depends on the degree to which **G** reflects the pleiotropic effects of new mutations [21-25]. Without a correlation between **G** and its mutational analogue, **M, G** is less likely to describe the nature of evolutionary constraints. Instead it will reflect other processes that change allele frequencies or the likelihood that alleles across multiple loci associate with one another, such as tight physical linkage or linkage disequilbrium [26], drift [27], correlated selection [23, 25, 28], and migration [22]. Additionally, little is known about the degree to which a genetic covariance is blind to pleiotropy that is “hidden” [29, 30]. Because **G** summarises the net contribution of all alleles affecting the genotype-to-phenotype (G-P) map, alleles with opposing contributions to genetic covariance effectively cancel each other out. Indeed, there are numerous G-P maps that could yield the same **G** [31, 32]. Given the central role of **G** in predicting evolutionary change, or lack thereof, it is important to understand its constituent alleles, their frequencies, and distribution of effects across multiple traits.

Historically, research programmes focussed on trait mapping have been isolated from those focussed on the estimation of **G** [31]. Although, multivariate mapping has gained popularity to leverage greater statistical power, studies that estimate multivariate effects in a manner useful to understanding **G** (or ***β*** and **Δ*Z***) remain rare. Work on *Mimulus gattatus* monkeyflowers has shown that allele frequency change at two multivariate QTL is predicted to cause substantial changes in **G** for life history traits [33]. Also, a multivariate GWAS of *Drosophila melanogaster* wing shape, showed how a multivariate approach provides a richer description of the pleiotropic nature of alleles that constitute the G-P map [34]. With regard to multivariate selection, a candidate gene association study on cuticular hydrocarbons (CHCs), a target of sexual selection in *Drosophila serrata*, estimated both vectors of multivariate SNP effect vectors alongside the selection gradient ***β*** allowing comparisons of their orientation in multi-trait space [35]. However, no study has yet jointly estimated how multivariate SNPs effects align with **G** and ***β***, both of which are required to predict multivariate responses to selection and uncover genetic constraints.

Here, we perform a genomic analysis of evolutionary constraints on a set of 7 cuticular hydrocarbon (CHC) traits in *D. serrata* that act as contact pheromones during mate choice [36-38]. Previous work indicates that mate choice by females generates strong sexual selection on male CHCs and that multivariate selection responses are constrained by comparatively low levels of multivariate genetic variance in **G** [16, 39]. Recent availability of whole-genome-sequenced lines for this system [40] provides an opportunity to expand the framework of Ivory-Church *et al*. [35] to a whole-genome view to investigate the alignment of SNP effects with not only the direction of selection, ***β***, but also the major axes of **G** that produce genetic constraints. Such frameworks may be useful as researchers increasingly access genomic data to study selection responses in wild populations.

## Methods

### Fly stocks

We used males from 91 lines from the *Drosophila serrata* Genome Reference Panel (DsGRP), which is a panel of 110 sequenced inbred lines derived from a single endemic population in South East Queensland, Australia [40]. Four males from each line were collected as virgins across two replicate rearing vials using light CO_2_ anaesthesia. For the mate choice assays, we used virgin male competitors and female ‘choosers’ from an inbred ‘tester’ line derived from a natural population in St Lucia, south-east Queensland, Australia approximately 5kms away from the population that founded the DsGRP. This inbred line carries a recessive mutation producing an orange-eye phenotype compared to the normal red-eye phenotype and allowed us to distinguish between focal and competitor males. Re-sequencing and variant calling of lines from the DsGRP has previously been described by Reddiex *et al*. [40]. In this study, SNPs were further filtered to be represented in at least 5% of the 91 lines. Highly corelated SNPs were removed using a sliding window approach implemented in PLINK v1.9 [41], with window size 100kb, step size of 20, and a r^2^ threshold of 0.5.

### Phenotyping mating success and CHCs

For each of the four males per line, a single focal male was placed in a vial with an orange-eyed male and an orange-eyed female. The vials were observed until either the focal male or the competitor successfully copulated with the female. At this point, the focal fly was assigned either a score of 1 if they were chosen by the female or a score of 0 if they were rejected. Upon mating, the focal male was immediately removed and had its CHCs extracted using a whole-body wash in 100μl of the solvent hexane. We used a standard gas chromatography method to quantify the amount of each CHC [36]. The areas under nine chromatograph peaks of interest (5,9-C_24:2_; 5,9-C_25:2_; 9-C_25:1_; 9-C_26:1_; 2-Me-C_26_; 5,9-C_27:2_; 2-Me-C_28_; 5,9-C_29:2_; 2-Me-C_30_) were measured and transformed into proportions to control for sample injected into the gas chromatograph machine as well as controlling any variation in the amount of CHCs extracted from the fly. To avoid the unit-sum constraints present for the 9 CHC ratios, we followed Aitchison [42] and generated log-contrasts for each of 8 CHC ratios by dividing the proportion of the CHC by the proportion of the remaining 9^th^ CHC, 9-C_26:1_, followed by taking the log of each of the 8 new variables. The log-contrast of 2-Me-C_30_ was removed from the analyses as it was being transferred between males and females during mating and became auto-correlated with mating success, a phenomenon that has been previous observed in this system [43], resulting in a total of 7 traits being analysed. Before statistical analysis we tested for multivariate outliers in males CHC expression by calculating the Mahalanobis distance using a p < 0.01 threshold based on a Chi-squared distribution [44]. Traits were standardised to have a mean of zero and a standard deviation of 1.

### Genomic estimation of G

We estimated the 7×7 additive genetic (co)variance matrix, **G**, for male CHC expression using a multivariate implementation of genome-based restricted maximum likelihood (GREML) [45] which is analogous to the animal model. Instead of fitting a relatedness matrix based on a pedigree we fitted a genomic relatedness matrix (GRM) based off the SNP data. Pairwise relatedness (*K*_*A*_) between lines (*j* and *k*) was estimated with the software GCTA [46] using the expression:

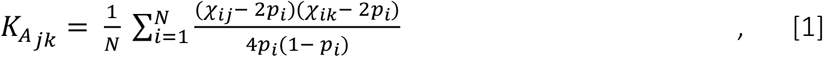

where, *χ*_*i,j*_ ∈ {0,2} is the number of copies of the reference allele for the *i* ^th^ SNP for inbred line *j* and *p*_*i*_ ∈ [0,1] is the population allele frequency. *N* is the total number of SNPs which was 1,652,276 for this analysis. In the denominator, we use the constant 4 instead of the commonly used 2 [47]. For populations of inbred lines, this produces a GRM with diagonal elements very close to 1 opposed to diagonal elements close to 2 (1 + the inbreeding coefficient) [48]). We implement this using the *--make-grm-inbred* option in GCTA.

Our multivariate linear model was specifically designed for populations where identical genotypes can be measured independently in multiple organisms such as inbred lines [49] and was performed using the R package, AsReml-R (VSN International). Here, the phenotypic values of replicate *r* of genotype *j* is modelled as:

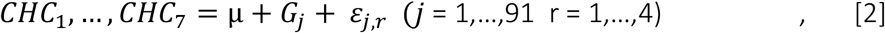

where *G* = (*G*_1_, …, *G*_7_) is the random effect of inbred line *j* and has a 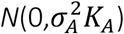 distribution. Errors *ε* = (*ε*_1_, …, *ε*_7_) are modelled with independent normal distributions with variances 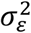. The number of independent dimensions of genetic variance that could be supported by the data was tested using factor-analytic modelling [50]. The linear model represented by Equation 2 was fit 7 times with the dimensionality of the **G** matrix constrained to be from 7 through to a single dimension. Then a series of nested likelihood ratio tests were performed to determine the number of significant dimensions [51]. The constraints on **G** were set using the reduced rank, rr(), option in AsReml-R.

### Estimating sexual selection

Standard multiple linear regression was used to estimate the vector of directional selection gradients ***β***. Before running the linear model, the binary measure of mating success was transformed to relative fitness by dividing all observations (0,1) by the mean mating success of the entire sample (i.e. total successes/total trials) Relative fitness was then modelled as the function of the intercept and the effects of each of the seven CHC traits as follows:

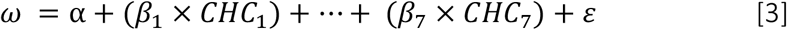

Equation3 was fit at the level of individuals using the R lm() function. For subsequent analyses, ***β*** was normalised to unit length.

### Multivariate genome-wide association analysis

We analysed the association between each SNP and CHC expression using the multivariate approach outlined in Equation 2 but with the addition of a fixed effect of *SNP* (*χ*_*i*_ ∈ {0,2}) using AsReml-R:

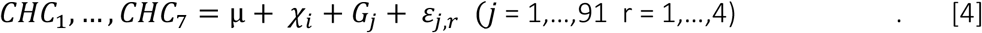

Typically, mixed model approaches to GWAS only estimate the polygenic random effect, modelled as the genomic relatedness matrix, once in a null model that excludes the fixed effect of SNP and then subsequent models including fixed SNP effects are then fit with the polygenic term variance components constrained to the null model estimates [46, 52, 53]. Our approach permits estimation of the polygenic random effect for each SNP, avoiding the double-fitting of SNPs common to popular GWAS software programs where SNPs are sequentially fit as a random effect (as a part of the GRM) and then as a fixed effect. Our approach slightly increases statistical power but comes at a cost of increased computation time [54, 55]. For each SNP, we extracted the estimated additive effect vectors, **s**, which were then used in subsequent analyses comparing them with major eigenvectors of **G** and ***β*** as outlined below.

### Genetic constraint

To assess genetic constraints on the response to selection, we calculated multiple statistics derived from the Lande equation using the selection gradient, ***β***, and the estimate of **G** from the model where all SNP effects are captured by the polygenic random effect but no individual SNP effects were fitted. First, we estimated the angle between the response to selection, **Δ*Z***, and ***β*** following Blows and Walsh [19] where:

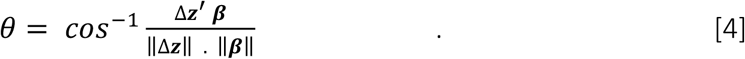

Second, we estimated the length of **Δ*Z*** that lies in the direction of ***β*** which is known as the evolvability, *e*(***β***), described by Hansen and Houle [20] as:

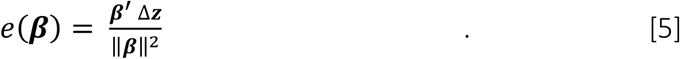

Finally, *ē*, the average evolvability of **G** was calculated using the R package ‘evolvability’ using a set of 10,000 randomly generated selection gradients [56].

### Orientation of SNP effects

For a fit of the GWAS model, a vector of SNP effects was generated, **s**. The individual elements of **s** describe the orientation of the SNP’s effect in multi-trait space and its length represents the effect size. To characterise the directions of **s**, we calculated a series of angles between **s** and the eigenvectors of **G** (θ_s,*g*1-7_) and between **s** and ***β*** (θ_s,***β***_) using analogous versions of Equation 4. Before the analysis, all vectors were normalised to have a length of one. Angles between two vectors were constrained to have values between 0° and 90°, that represent perfect alignment or misalignment respectively.

We considered three methods to develop null expectations for these orientation parameters to determine whether the angles we observed for significant SNPs deviate from what would be expected by chance. First, which we ultimately rejected, was to randomly draw 10,000 vectors uniformly distributed in a k-dimensional hypersphere and use the angles between these vectors and the eigenvectors of **G** and ***β*** as a null distribution. In our 7-dimensional case, the median angle between two random vectors under this sampling strategy is approximately 73°. However, this assumes **G** is spherical and therefore that correlations between all traits are zero, which is not representative of randomly generated matrices in the presence of sampling variance [57, 58]. Random matrix theory predicts a pattern of decreasing eigenvalues in estimated data even in the absence of any biological signal of covariance among traits [57, 58]. What this means is that angles between SNP vectors, **s**, and the eigenvectors of **G** will be expected to also show a pattern of decaying alignment from the first through to the last eigenvector of **G**. Thus, we needed to account for this phenomenon and assess whether decay observed deviates from the null expectation. Second, we took 1,000 stratified samples of non-significant MAF-matched SNPs using the R package *sampling*. Each sample had the same number of SNPs, and retained the same distribution of minor allele frequencies, as the significant SNPs. For each sample, we took the median value for each orientation parameter to build the null distribution. Third, we generated samples of 10,000 random 7 element vectors drawn from a multivariate normal distribution where the variance-covariance matrix was **G** estimated from Equation 2. Similar to Pitchers *et al*. [34], we found that the second and third methods produce equivalent null distributions and chose to use the null distribution derived from the stratified samples of non-significant SNPs.

## Results

### Genome-based REML estimates of the G matrix

We tested for the number of significant genetic dimensions of genetic variance supported by the GREML model using factor analytic modelling. There was a significant worsening in model fit when testing a model with five dimensions against a model with four dimensions (LRT: *χ*^2^ = 58.93, d.f. = 3, p = 9.9 × 10^−13^). This result indicated we had statistical support for 5 out of a possible 7 dimensions, a result also underscored by the AIC values, for which the 5-dimension model was lowest (Table 1).

**Table 1.** Summary of linear model fits for factor analytic modelling to test the dimensionality of the genetic variance- covariance matrix, **G**_SNP_, estimated from the genomic relatedness matrix.

| Dimensions<br>fitted | Model Fit |  |  |
| --- | --- | --- | --- |
|  | -2LL | AIC | no.<br>parameters |
| 7 | 207.49 | 319.49 | 56 |
| 6 | 207.51 | 317.51 | 55 |
| **5** | **211.47** | **317.47** | **53** |
| 4 | 270.40 | 370.40 | 50 |
| 3 | 394.93 | 486.93 | 46 |
| 2 | 558.62 | 640.62 | 41 |
| 1 | 763.01 | 833.01 | 35 |

### Sexual selection and genetic constraint

We found significant sexual selection on male CHC expression (ANOVA: F_7,347_ = 7.01, p = 6.37×10^−8^), where CHC expression explained 11% of the variance in relative fitness, a level typical of other studies in this system [43]. Individually, three of the seven CHCs had significant main effects on relative fitness (t-Test: 2MeC_26_: t = 4.78, p = 2.62×10^−6^; 2MeC_28_: t = -2.75, p = 6.26×10^−3^; 5,9C_29:2_: t = 5.94, p = 6.82×10^−9^). Two of the seven CHC are under positive selection where the other CHCs are subject to negative sexual selection (Table 2).

**Table 2:** The additive genetic (co)variance matrix **G**_SNP_ and sexual selection gradient ***β*** for male *Drosophila serrata* from the DsGRP. Elements of the diagonal and lower triangle of the matrix are trait variances and covariances while the upper triangle contains trait correlations. Asterisks represent significant partial regression coefficients at p < 0.05 from multiple regression analysis. The matrix shown is from a model assuming an unstructured variance-covariance matrix (i.e. AsReml-R us()) for the additive genetic effect. Note that estimates need to be halved to approximate the V_A_ present in the *outbred base population* under an assumption of additivity ref [48] p.265.

| | 5,9-C<sub>24:2</sub> | 5,9-C<sub>25:2</sub> | 9-C<sub>25:1</sub> | 2-MeC<sub>26</sub> | 5,9-C<sub>27:2</sub> | 2-MeC<sub>28</sub> | 5,9-C<sub>29:2</sub> | $\boldsymbol{\beta}$ |
| --- | --- | --- | --- | --- | --- | --- | --- | --- |
| 5,9-C<sub>24:2</sub> | **0.382** | 0.926 | 0.318 | 0.567 | 0.087 | 0.548 | 0.179 | -0.069 |
| 5,9-C<sub>25:2</sub> | **0.381** | **0.444** | 0.444 | 0.434 | 0.317 | 0.593 | 0.353 | -0.222 |
| 9-C<sub>25:1</sub> | **0.127** | 0.191 | **0.416** | 0.050 | 0.156 | 0.215 | 0.029 | -0.052 |
| 2-MeC<sub>26</sub> | **0.214** | 0.177 | 0.020 | **0.373** | 0.110 | 0.615 | 0.422 | 0.608* |
| 5,9-C<sub>27:2</sub> | **0.039** | 0.156 | 0.074 | 0.049 | **0.543** | 0.488 | 0.678 | -0.113 |
| 2-MeC<sub>28</sub> | **0.214** | 0.250 | 0.088 | 0.238 | 0.227 | **0.400** | 0.826 | -0.396* |
| 5,9-C<sub>29:2</sub> | **0.040** | **0.085** | **0.007** | **0.093** | **0.180** | **0.188** | **0.130** | 0.635* |

We found that the pattern of genetic (co)variance, represented by **G**, rotated the response to selection away from the direction defined by ***β*** where the angle between **Δ*Z*** and ***β***, *θ*_**Δ*Z***,***β***_= 51° (Figure 1). We estimated the degree to which the response to selection occurred in the direction of ***β***, i.e. the evolvabilty, to be *e*(***β***) = 0.126 (Figure 1), which is considerably less (about one third) than the average evolvability of **G**_SNP_ for this population, ***ē*** = 0.387. Together these results suggest that the response to directional selection is indeed constrained by the multivariate distribution of genetic variance in this population.

**Figure 1:**
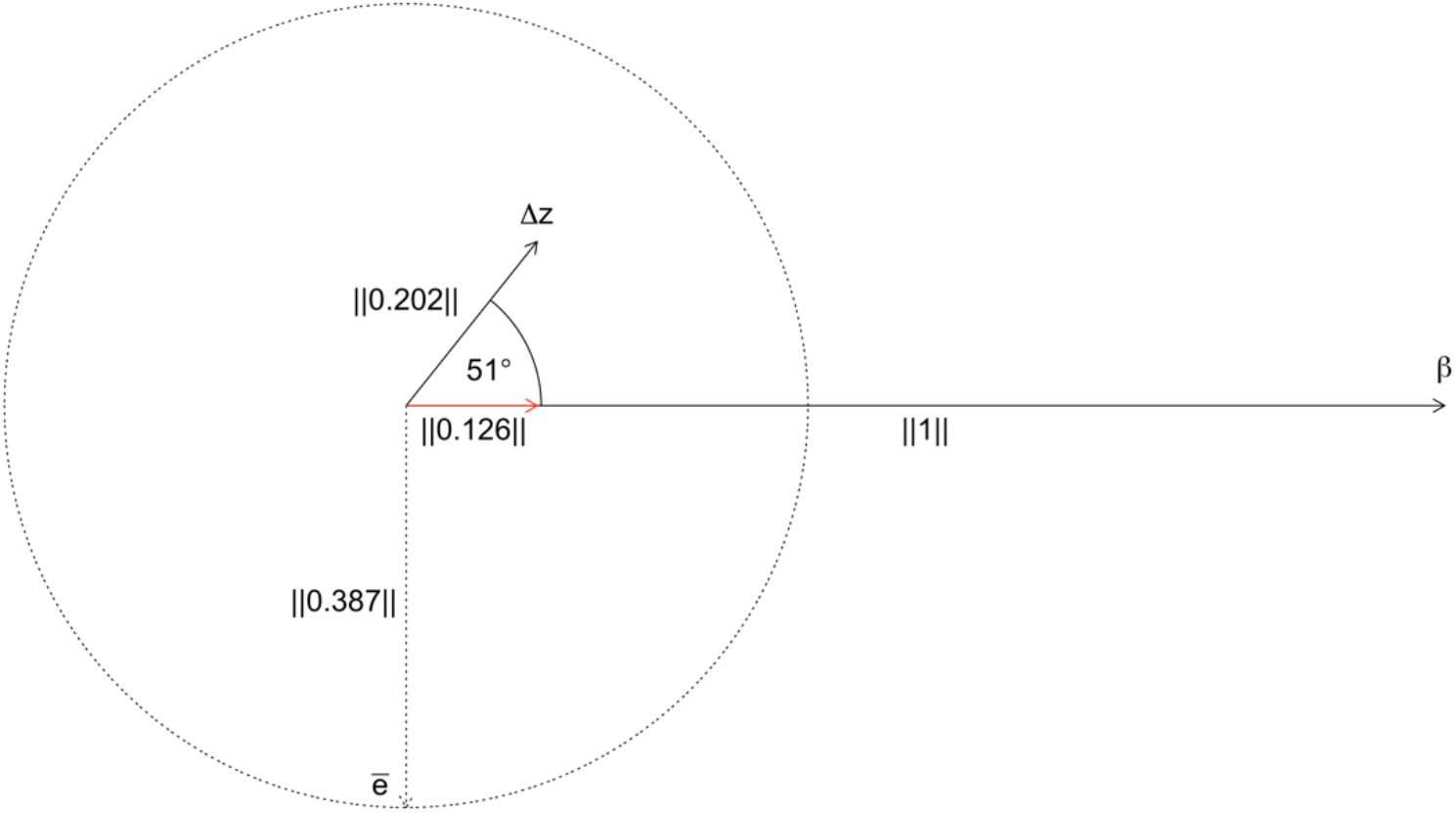
Genetic constraint on the response to selection on male CHCs in *D*.*serrata*. Vector correlation between the response to selection, as calculated from the Lande equation, and ***β, θ***_**Δ*Z***,***β***_. Also shown is the evolvability ***e***(***β***) depicted as the red arrow and average evolvability ***ē*** shown as the radius of the dotted circle.

### Multi-trait GWAS

Of the 1,652,276 autosomal SNPs analysed, 532 had significant multivariate associations with CHC variation (FDR <5%, [59]; Figure 2). The Q-Q plot of the observed p-value distribution indicated inflation against expected values (Fig S1); a signal of genomic inflation. Several factors can contribute towards genomic inflation in GWAS of polygenic traits [60]. While uncontrolled relatedness between genotypes is often of principal concern, the DsGRP population has extremely low signals of relatedness and for the lines we have considered here, there was no significant relatedness among genotypes [61]. Linkage disequilibrium also contributes to genomic inflation as SNPs in close proximity to causal SNPs will have elevated test statistics [60]. Here, the effect of linkage disequilibrium on genomic inflation should be minimal, as we used LD pruning to remove highly correlated SNPs that are close to one another before conducting GWAS. However, low frequency SNPs that are far apart along a chromosome, or even on different chromosomes, may still be highly correlated in studies with low sample sizes and many SNPs such as ours [62]. Houle and Márquez [63] refer to this phenomenon as ‘rarity disequilibrium’ as it arises due to the relatively small number of permutations of individuals that can possess a rare SNP. Rarity disequilibrium may explain a portion of the genomic inflation in this study as it is consistent with the higher observed inflation of low frequency SNPs compared to common SNPs (Fig S1). However, within the 532 statistically significant SNPs, we found little evidence for this type of long-range LD. Out of the possible 141,246 pair-wise combinations of significant SNPs, only 48 pairs had an r^2^ > 0.5 (Fig S2). Finally, highly polygenic traits will display higher values for genomic inflation compared to traits with a simpler genetic architecture, reflecting true, but small, effects. Consistent with this idea, we analysed of the p-value distribution using the approach of Storey and Tibshirani [64] and estimated *η*_0_ = 0.8832, which suggest that 88% of the SNPs had no phenotypic effect.

**Figure 2:**
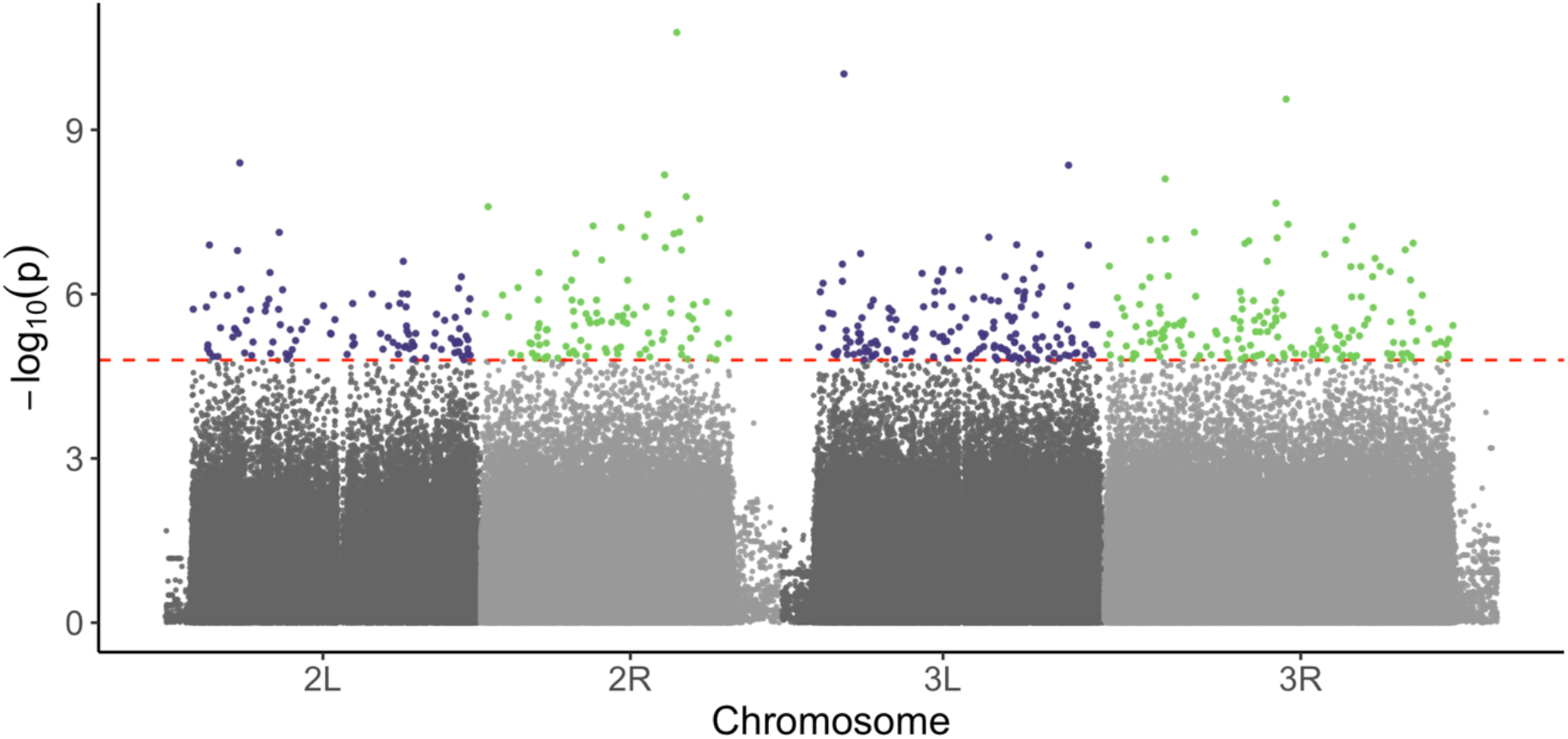
Manhattan plot for the multivariate GWAS model fitted to seven CHC traits in *D. serrata*. Horizontal line corresponds to the FDR threshold of 5% [59]. The appearance of telomeric regions is due to their much lower (approximately 100-fold) SNP density than other genomic regions. 38,887 SNPs analysed on unplaced but confirmed-autosomal scaffolds are not shown.

### Orientation of SNP effects with major axes of the G matrix

We compared the orientation between the vectors of significant SNP effects, **s**, with the eigenvectors of **G** by calculating the angles between them. To determine whether SNP vector alignments were better or worse than expected by chance, we generated 1000 random samples of 532 SNPs from the non-significant SNP vectors while matching the distribution of minor allele frequencies. When the median angle between our observed set of 532 SNPs and our null samples was in the upper or lower 2.5% percentiles of the distribution of random samples we declared the result significant.

The first eigenvector of **G**, ***g***_max_ comprised 50% of the genetic variance and was characterised by loadings of a common sign and of quite similar magnitude (Table 3), a result consistent with previous studies of the *D. serrata* CHC **G** matrix [16, 39, 43]. The observed median angle between the set of significant SNP vectors and this leading eigenvector, *θ*_*snp,gmax*,_was 60°. When compared to the null distribution we see that the observed median angle was approximately 6° larger than expected by chance and no overlap between the median values of the 1000 532-SNP subsamples and our observed value (Figure 3). Thus, there was significant misalignment between ***g***_max_ and significant SNP effects.

**Table 3:**
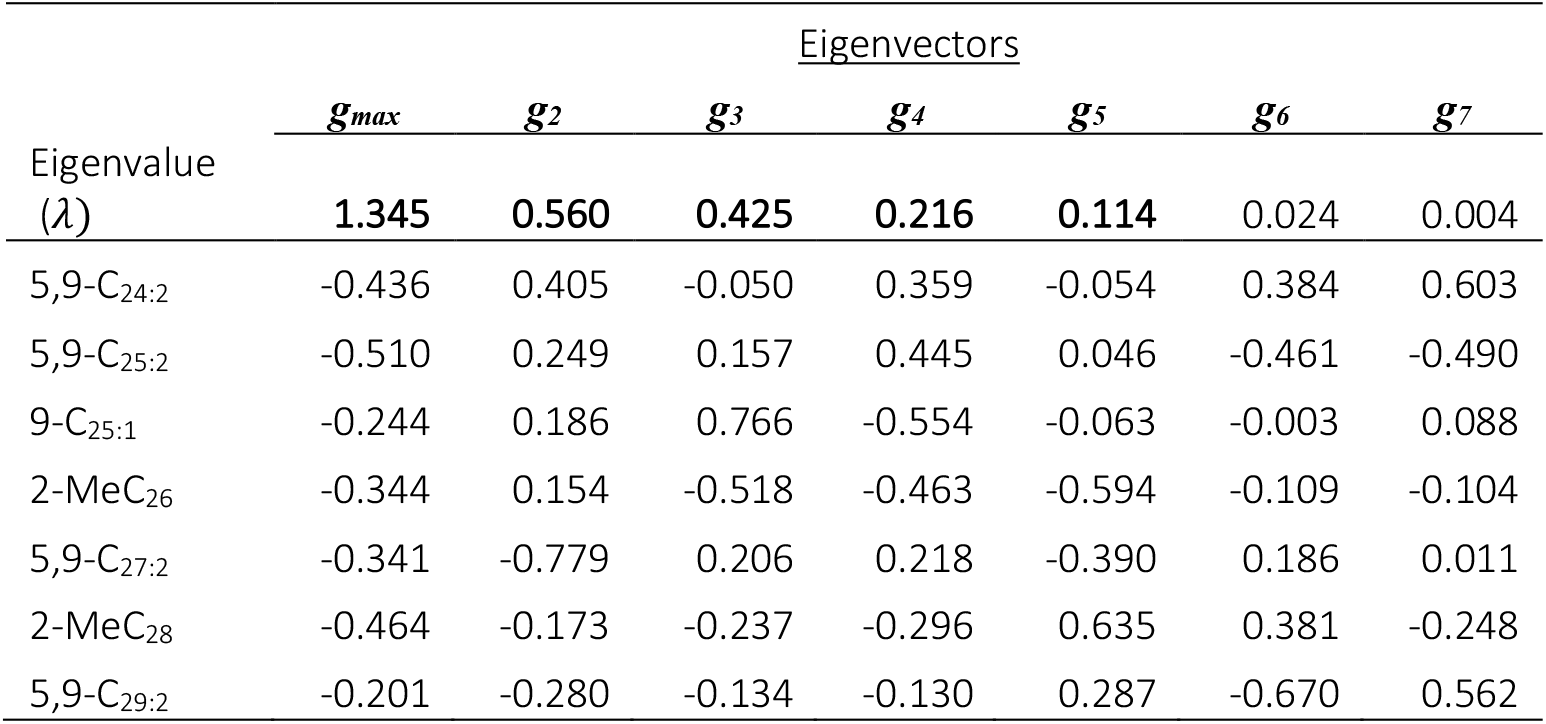
Eigenvectors and eigenvalues of G_SNP_ for seven CHC traits of *D. serrata*. Eigenvalues in bold reflect the vectors within phenotypic trait space with statistical support via factor analytic modelling (Table 1). The vectors shows are from a model assuming an unstructured variance-covariance matrix (i.e. ASREML-R us()) for the additive genetic effect.

**Figure 3:**
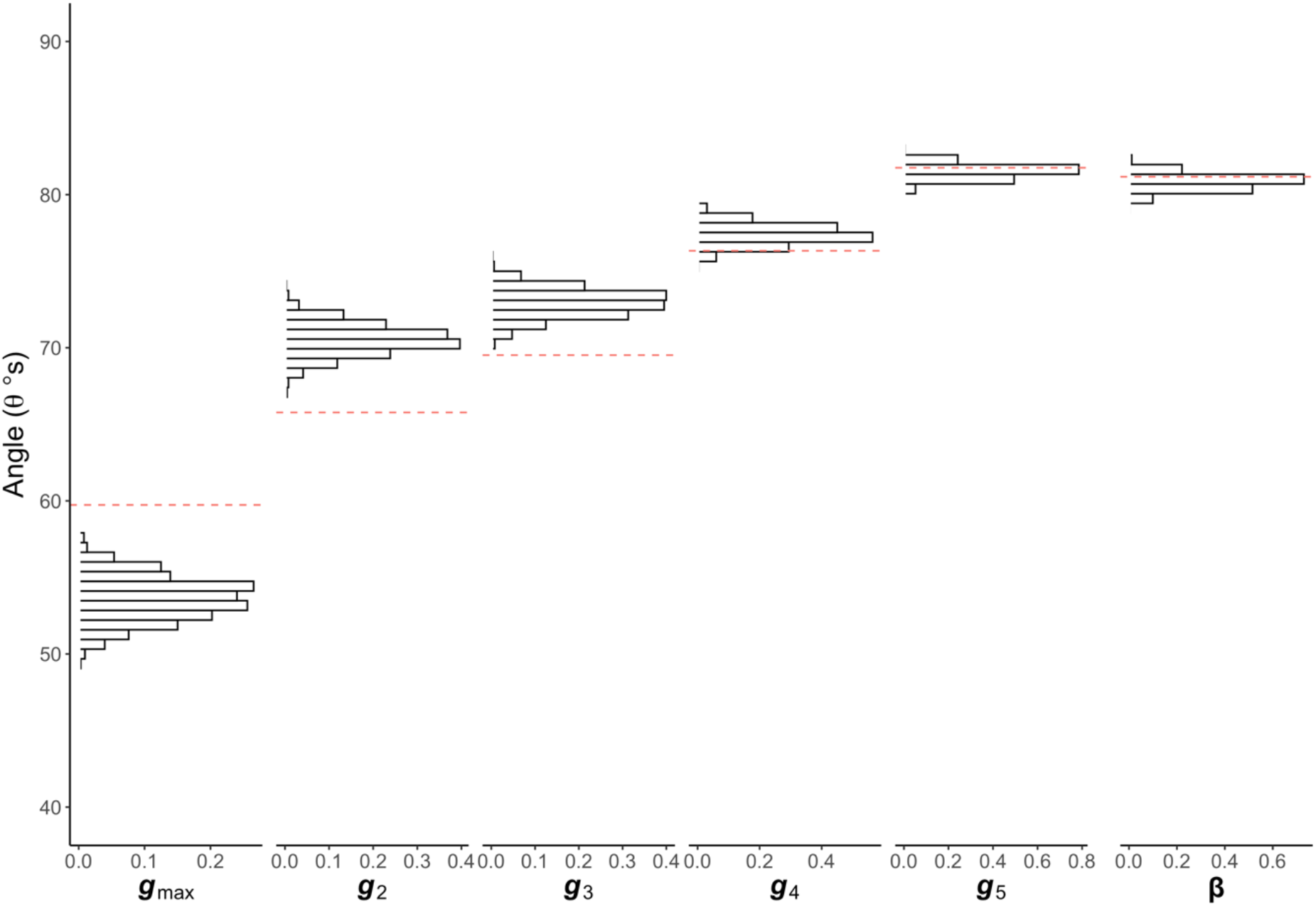
Median vector correlations between the first 5 eigenvectors of **G** and the vector of SNP effects. Red dashed lines represent the median angle (in degrees) between the SNP vector and each eigenvector of **G** observed in significant SNPs. Histograms represent a null distribution of expected median angles derived from 1000 random samples, without replacement, of 532 non-significant SNPs (the same number as significant SNPs).

The second eigenvector of **G, *g***_2_ comprised 21 % of the total genetic variance and had loadings reflecting a contrast between long- and short-chained CHCs with opposing signs between CHCs above and below 27 carbon atoms (Table 3). In contrast to the result for ***g***_max_, estimates of SNP vectors had alignments closer to this eigenvector than expected by chance. The observed median angle, *θ*_*snp,g*2_, was 65.8° whereas the null median was 70.5° and again the distribution did not overlap with the observed angle (Figure 3). We saw a similar signal of better than expected alignment for the third eigenvector of **G, *g***_3_ which explained 16 % of the total genetic variance and had loadings that to a large extent, contrasted the both methyl-branched alkanes 2-MeC_26_ and 2-MeC_28_ with the monoene 9-C_27:1_. The observed angle *θ*_*snp,g*3_ of 69.5° did not overlap with any of the null samples for which the median orientation was approximately 73° (Figure 3). In contrast, the situation changed for the smaller eigenvectors ***g***_4_ and ***g***_5_ for which there was no significant signal of alignment or misalignment and in all cases observed angles overlapped the null distributions (Figure 3). Because factor analytic modelling indicated that genetic variance could not be supported by the data for a 6^th^ and 7^th^ dimension, the angles between SNP effects and ***g***_6_ and ***g***_7_ were not analysed further.

### Orientation of SNP effects with directional selection

As discussed above, the structure of **G** biased the predicted response to sexual selection, **Δ*Z***, 51° away from the direction of selection on male CHCs, ***β***. Statistically significant SNPs were generally poorly aligned to ***β***, with a median angle of *θ*_*snp*,***β***_ = 81.2° and only two of the 532 significant SNPs had *θ*_*snp*,***β***_ < 45°. Our observed median angle of 81.2°, however largely follows the null expectation for a variance-covariance matrix shaped like our estimated **G** and sits in the middle of the null distribution (Figure 3). This suggests that segregating variants affecting male CHC variation are no better or worse aligned with the linear combination of CHCs preferred by females than expected.

## Discussion

A major goal of evolutionary genetics is to understand the direction and rate of evolution given that multiple traits experience selection simultaneously but often share genetic architecture [11]. Historically, the analysis of the genetic (co)variance matrix **G** and the mapping of genetic variants to phenotypic variation have occurred in isolation due to the fundamental incompatibilities in experimental design [31]. Consequently, the investigation of how pleiotropy constrains phenotypic evolution has in large part relied on estimates of **G**, and its spectral decomposition into eigenvectors and eigenvalues. The use of **G** to infer genetic constraints has been sometimes criticised by claims that eigenvectors of **G** don’t necessarily reflect the pleiotropic nature of the G-P map [32, 65, 66]. Chebib and Guillaume [25] have recently shown that the utility of **G** for evolutionary inference, especially over the longer term, relies on a correlation between **G** and its mutational analogue, **M**. While mutational pleiotropy can be assessed in model organisms through the estimation of genetic covariance in mutation accumulation studies [67-69], such experiments are not feasible for free-living populations of non-model species, the study of which is critical to understand phenotypic evolution in the wild. However, mapping in populations where **G** and multivariate allelic effects can be jointly estimated could potentially bridge this divide. Here we have identified common SNPs that underlie a G-P map and characterised their orientation in multi-trait space in relation to the eigenvectors of **G**, and multivariate directional selection, ***β***. Our approach permits investigation of the extent to which these key evolutionary parameters for predicting responses to selection reflect the underlying G-P map.

### Alignment of the genotype-phenotype map and the major axes of genetic variation

We assessed the alignment of SNP effects with the major axes of genetic (co)variance using vector correlations. What we find interesting is that there was statistical support for both significant alignment and misalignment depending upon the specific eigenvector being examined. SNPs were more closely aligned to the second and third eigenvectors of **G, *g***_2_ and ***g***_3_, than expected by chance (Fig 3). Stated another way, there was an enrichment of SNPs that affect CHCs in the direction of ***g***_2_ and ***g***_3_ that passed our statistical significance threshold. Closer than average alignment for ***g***_2_ and ***g***_3_ is to some extent consistent with our current understanding of the biosynthetic pathways of CHC development. For example, the loadings of ***g***_2_ show a contrast between long- and short-chained CHCs with opposing signs between CHCs above and below 27 carbon atoms. Genes such as elongases have been shown to have phenotypic effects on carbon chain length in CHCs in flies [70] and reductases affect fatty alcohol-based pheromone chain length in other insects [71]. Moreover, the three strongest loading traits on ***g***_3_ highlight a contrast between strong positive effects on the only monoene analysed, 9-C_25:1_, and negative effects on the two methyl-branched alkanes, 2-MeC_26_ and 2-MeC_28_. The biosynthesis of methyl-branched alkanes is known to require a different class of fatty acid synthase (FAS) gene to those required for the synthesis of monenes, dienes and other alkanes [72].

The phenotypic effects represented by the major axis of genetic variance, ***g***_max_, are different and involve loadings of a common sign and similar magnitude suggesting pleiotropic alleles that affect CHCs in similar ways. Our observation of poor alignment of SNP effects with ***g***_max_ could be explained by two scenarios. First, mutation may seldom generate phenotypic effects aligned with ***g***_max_ and instead ***g***_max_ reflects an average of many varied directions of effect. Second, if mutations do commonly align with ***g***_max_, any well-aligned mutations may be more effectively removed from the population by selection or driven to lower frequencies, leaving us with the poorer aligned SNPs segregating at higher frequencies making them detectable by association mapping. It is noteworthy that CHC ***g***_max_ has been shown to experience strong stabilising selection, at the genetic as opposed to phenotypic level, in *D. serrata* [73, 74]. This observation is more consistent with the scenario of our mapped alleles being a subset of the variation introduced by mutation. An interesting question is whether any variants well aligned with ***g***_max_ tend to have very small effect sizes making them difficult to detect or whether they have large effects but are rare. Distinguishing between these alternatives is likely to require much larger samples sizes than used in the present study.

### The alignment of directional selection and the G-P map

We found limited potential for male CHCs to evolve in the direction that would maximise their attractiveness to females. Due to the structure of **G**, the response to selection (**Δ*Z***) is predicted to be rotated 51° away from the direction of ***β***, furthermore, the estimates of evolvability (*e*(***β***)) and average evolvability (*ē*) shows a 67% reduction in the response to selection in the direction of ***β*** compared to the average evolutionary potential of **G**. This high genetic constraint is consistent with previous studies of CHC expression in *Drosophila serrata* [43, 51, 75], and other studies that provide strong evidence of multivariate genetic constraint in other organisms [76-79] but see [80]. The apparent lack of genetic variance in the direction of sexual selection is reflected in the fact that most SNP effects were poorly aligned with ***β*** (Figure 3). This may reflect a developmental constraint where single genes involved in CHC production cannot mechanistically produce CHCs that maximise attractiveness. Thus ***β*** may be highly polygenic, and reflective of the effect across many different types of loci.

Alternatively, and similar to the argument for ***g***_max_, any variants that are closely aligned with ***β***, may occur at very low/high frequencies at which GWAS are no longer suitable to detect them. Interestingly, we observed bias in the frequency of the ***β*** increasing allele. Alleles increasing ***β*** and therefore male attractiveness were more often the minor frequency allele than expected against 1000 random draws of 532 MAF-matched control SNPs (χ^2^ = 19.789, df = 1, p = 8.65e-06). This finding is consistent with the idea that there may be a fitness cost associated with high ***β*** expression in this species, an idea with some experimental support from artificial selection studies conducted on CHC ***β*** in this species [81, 82]. Future studies may be able to address these questions by experimentally manipulating population allele frequencies either through crossing selected lines or by artificially applying (or removing) selection alongside SNP genotyping.

## Caveats and Conclusion

Our study used multivariate GWAS to ascertain SNPs with the largest effects and describe the geometry of their effects. It is important to highlight that the ascertainment of SNPs and the geometry of **G** itself are not independent. The eigenvectors of **G** describe the directions of trait-space with different amounts of genetics variance, and because genetic variance is a major determinant of the statistical power of a GWAS, there is an inherent bias towards finding SNPs that are most closely aligned to ***g***_max_ compared to the other eigenvectors of **G**. This is why it is essential to calibrate orientation results against a null distribution. An additional improvement to the approach outlined here would be to collect multiple datasets from the same population so that **G** and multivariate SNP effects could be estimated independently.

The utility of **G** to predict evolutionary response of multiple phenotypes beyond the short-term remains an active topic of research. As discussed above, for **G** to accurately predict longer-term evolutionary responses, there must be a meaningful relationship between itself and the pleiotropic effects of new mutations [25]. For the study of natural populations, this poses a considerable challenge as researchers do not have the capacity to remove natural selection or to standardise the genetic background of organisms that allows mutational genetic (co)variance to be estimated for laboratory populations. Here, we use GWAS as an alternative framework to investigate the relationship between **G** and the pleiotropic effects of mutation, albeit for mutations that are segregating at common frequencies in the population. We found that the pleiotropic effects of SNPs that underlie the G-P map significantly deviate from what is expected given the geometry of **G**, which may alter how we anticipate how these traits may evolve in the future. As genome-wide sequence data is rapidly becoming available for wild populations [83, 84], we think that frameworks like that presented here, where evolutionary parameters such as **G, *β*** and the G-P map are jointly estimated, can improve our understanding of evolutionary constraints and selection responses.

## Acknowledgements

This work was supported by funding from The Australian Research Council and The University of Queensland. AR was supported by a Commonwealth RTP scholarship. We thank M. Hall for comments on an earlier draft of the manuscript and S. Allen for assistance with the *D. serrata* genome and creating Figure 2. N. Appleton, S. Allen, and T.P. Gosden assisted with fly rearing, CHC extraction and processing the mate choice trials.

## Supplementary Figures

**Figure S1:**
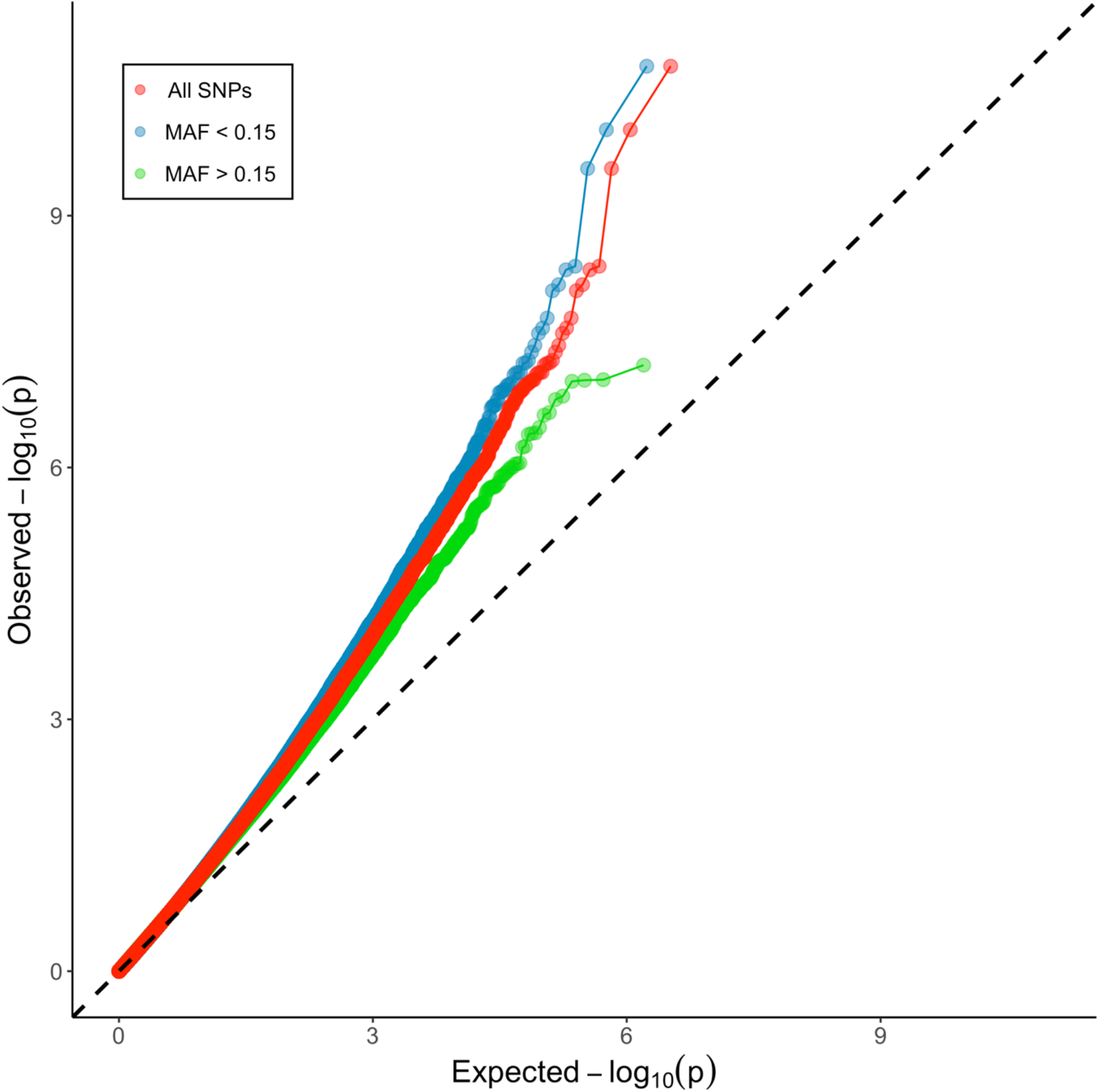
Q-Q Plot of observed vs expected -log10(p-value) distributions from the 1,652,276 single SNP multivariate GWAS mixed effects models. Series are plotted for SNPS with different minor allele frequencies.

**Figure S2:**
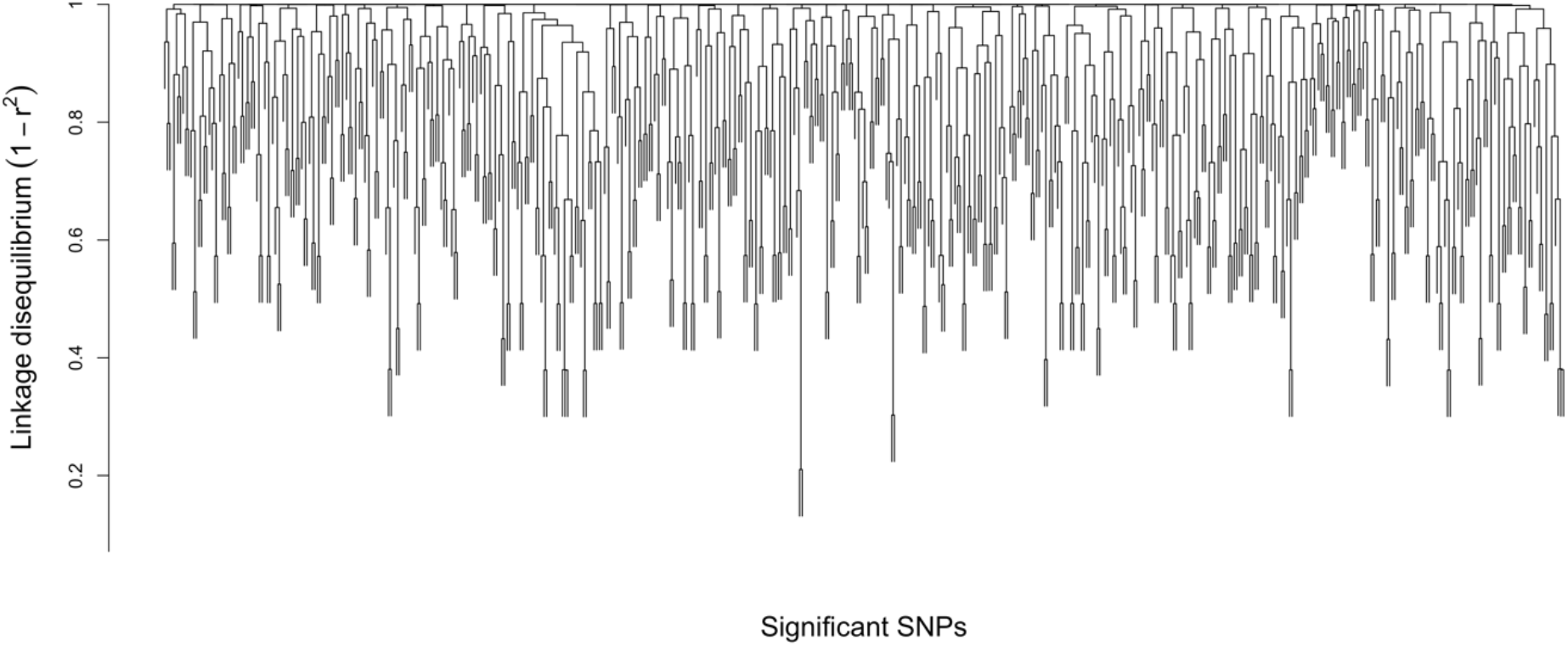
Dendrogram of linkage disequilibrium of the 532 significant SNPs. Pair-wise linkage disequilibrium between significant SNPs was calculated as the squared correlation of inter-variate allele counts using PLINK. The resulting 532×532 correlation matrix was converted to a distance matrix before being subject to hierarchical clustering using the ‘*hclust’* function in R.

## References

[1] Mousseau, T.A. & Roff, D.A. 1987 Natural-selection and the heritability of fitness components. Heredity 59, 181–197.

[2] Lynch, M. & Walsh, B. 1998 Genetics and Analysis of Quantitative Traits. Sunderland, Sinauer Associates.

[3] Hill, W.G. & Caballero, A. 1992 Artificial selection experiments. Annual Review of Ecology and Systematics 23, 287–310. (doi:10.1146/annurev.es.23.110192.001443.)

[4] Kingsolver, J.G., Hoekstra, H.E., Hoekstra, J.M., Berrigan, D., Vignieri, S.N., Hill, C.E., Hoang, A., Gibert, P. & Beerli, P. 2001 The strength of phenotypic selection in natural populations. Am. Nat. 157, 245–261.

[5] Hereford, J., Hansen, T.F. & Houle, D. 2004 Comparing strengths of directional selection: How strong is strong? Evolution 58, 2133–2143.

[6] Johnson, T. & Barton, N. 2005 Theoretical models of selection and mutation on quantitative traits. Philosophical Transactions of the Royal Society of London. Series B - Biological Sciences 360, 1411–1425. (doi:10.1098/rstb.2005.1667.)

[7] Fisher, R.A. 1930 The genetical theory of natural selection. Oxford, UK, The Clarendon Press.

[8] Houle, D. 1992 Comparing evolvability and variability of quantitative traits. Genetics 130, 195–204.

[9] Merila, J., Sheldon, B.C. & Kruuk, L.E. 2001 Explaining stasis: microevolutionary studies in natural populations. Genetica 112, 199–222.

[10] Kruuk, E.B., Slate, J., Pemberton, J.M., Brotherstone, S., Guinness, F. & Clutton-Brock, T. 2002 Antler size in red deer: heritability and selection but no evolution. Evolution 56, 1683–1695.

[11] Walsh, B. & Blows, M.W. 2009 Abundant genetic variation + strong selection = multivariate genetic constraints: a geometric view of adaptation. Annual Review of Ecology, Evolution, and Systematics 40, 41–59. (doi:10.1146/annurev.ecolsys.110308.120232.)

[12] Lande, R. 1979 Quantitative genetic analysis of multivariate evolution, applied to brain: body size allometry. Evolution 33, 402–416. (doi:10.2307/2407630.)

[13] Kirkpatrick, M. 2009 Patterns of quantitative genetic variation in multiple dimensions. Genetica 136, 271–284. (doi:10.1007/s10709-008-9302-6.)

[14] Blows, M.W. & McGuigan, K. 2014 The distribution of genetic variance across phenotypic space and the response to selection. Molecular Ecology 24, 2056–2072. (doi:10.1111/mec.13023.)

[15] Gomulkiewicz, R. & Houle, D. 2009 Demographic and genetic constraints on evolution. American Naturalist 174, E218–229. (doi:10.1086/645086.)

[16] Chenoweth, Stephen F., Rundle, Howard D. & Blows, Mark W. 2010 The contribution of selection and genetic constraints to phenotypic divergence. The American Naturalist 175, 186–196. (doi:10.1086/649594).

[17] McGuigan, K., Chenoweth, S.F. & Blows, M.W. 2005 Phenotypic divergence along lines of genetic variance. Am Nat 165, 32–43.

[18] Lande, R. & Arnold, S.J. 1983 The measurement of selection on correlated characters. Evolution 37, 1210–1226.

[19] Blows, M. & Walsh, B. 2009 Spherical cows grazing in Flatland: constraints to selection and adaptation. In Adaptation and Fitness in Animal Populations: Evolutionary and Breeding Perspectives on Genetic Resource Management (eds. J. van der Werf, H.-U. Graser, R. Frankham & C. Gondro), pp. 83–101. Dordrecht, Springer Netherlands.

[20] Hansen, T.F. & Houle, D. 2008 Measuring and comparing evolvability and constraint in multivariate characters. Journal of evolutionary biology 21, 1201–1219. (doi:10.1111/j.1420-9101.2008.01573.x.)

[21] Arnold, S.J., Bürger, R., Hohenlohe, P.A., Ajie, B.C. & Jones, A.G. 2008 Understanding the evolution and stabilty of the G-matrix. Evolution 62, 2451–2461. (doi:10.1111/j.1558-5646.2008.00472.x.)

[22] Guillaume, F. & Whitlock, M.C. 2007 Effects of migration on the genetic covariance matrix. Evolution 61, 2398–2409.

[23] Jones, A.G., Arnold, S.J. & Bürger, R. 2003 Stabilty of the G-matrix in a population experiencing pleiotropic mutation, stabilizing selection, and genetic drift Evolution 57, 1747–1760. (doi:10.1111/j.0014-3820.2003.tb00583.x.)

[24] Steppan, S.J., Phillips, P.C. & Houle, D. 2002 Comparative quantitative genetics: evolution of the G matrix. Trends in ecology & evolution 17, 320–327. (doi:https://doi.org/10.1016/S0169-5347(02)02505-3.)

[25] Chebib, J. & Guillaume, F. 2017 What affects the predictability of evolutionary constraints using a G-matrix? The relative effects of modular pleiotropy and mutational correlation. Evolution 71, 2298–2312. (doi:10.1111/evo.13320.)

[26] Lande, R. 1980 The Genetic Covariance between Characters Maintained by Pleiotropic Mutations. Genetics 94, 203–215.

[27] Griswold, C.K., Logsdon, B. & Gomulkiewicz, R. 2007 Neutral evolution of multiple quantitative characters: a genealogical approach. Genetics 176, 455–466. (doi:10.1534/genetics.106.069658.)

[28] Jones, A.G., Arnold, S.J. & Bürger, R. 2007 The mutation matrix and the evoution of evolvability. Evolution 61, 727–745. (doi:10.1111/j.1558-5646.2007.00071.x.)

[29] Wagner, G.P. 1989 Multivariate Mutation-Selection Balance with Constrained Pleiotropic Effects. Genetics 122, 223–234.

[30] Baatz, M. & Wagner, G.P. 1997 Adaptive inertia caused by hidden pleiotropic effects Theoretical Population Biology 51, 49–66.

[31] Kelly, J.K. 2009 Connecting QTLS to the g-matrix of evolutionary quantitative genetics. Evolution 63, 813–825. (doi:10.1111/j.1558-5646.2008.00590.x.)

[32] Gromko, M.H. 1995 Unpredictability of the correlated response to selection: pleiotropy and sampling interact. Evolution 49, 685–693. (doi:10.1111/j.1558-5646.1995.tb02305.x.)

[33] Scoville, A., Lee, Y.W., Willis, J.H. & Kelly, J.K. 2009 Contribution of chromosomal polymorphisms to the G-matrix of Mimulus guttatus. New Phytologist 183, 803–815. (doi:10.1111/j.1469-8137.2009.02947.x.)

[34] Pitchers, W., Nye, J., Márquez, E.J., Kowalski, A., Dworkin, I. & Houle, D. 2019 A multivariate genome-wide association study of wing shape in Drosophila melanogaster. Genetics 211, 1429. (doi:10.1534/genetics.118.301342.)

[35] Ivory-Church, J., Frentiu, F.D. & Chenoweth, S.F. 2015 Polymorphisms in a desaturase 2 ortholog associate with cuticular hydrocarbon and male mating success variation in a natural population of Drosophila serrata. J. Evol. Biol. 28, 1600–1609. (doi:10.1111/jeb.12679.)

[36] Blows, M.W. & Allan, R.A. 1998 Levels of mate recognition within and between two Drosophila species and their hybrids. Am. Nat. 152, 826–837.

[37] Higgie, M., Chenoweth, S. & Blows, M.W. 2000 Natural selection and the reinforcement of mate recognition. Science (New York, N.Y.) 290, 519–521.

[38] Chenoweth, S.F. & Blows, M.W. 2005 Contrasting mutual sexual selection on homologous signal traits in Drosophila serrata. American Naturalist 165, 281–289. (doi:10.1086/427271.)

[39] Hine, E., Chenoweth, S.F. & Blows, M.W. 2004 Multivariate quantitative genetics and the lek paradox: genetic variance in male sexually selected traits of Drosophila serrata under field conditions Evolution 58, 2754–2762. (doi:https://doi.org/10.1111/j.0014-3820.2004.tb01627.x.)

[40] Reddiex, A.J., Allen, S.L. & Chenoweth, S.F. 2018 A Genomic Reference Panel for Drosophila serrata. G3: Genes|Genomes|Genetics 8, 1335. (doi:10.1534/g3.117.300487.)

[41] Purcell, S., Neale, B., Todd-Brown, K., Thomas, L., Ferreira, M.A.R., Bender, D., Maller, J., Sklar, P., de Bakker, P.I.W., Daly, M.J., et al. 2007 PLINK: a tool set for whole-genome association and population-based linkage analyses. Am. J. Hum. Genet. 81, 559–575. (doi:10.1086/519795.)

[42] Aitchison, J. 1986 The Statistical Analysis of Compositional Data. London, Chapman and Hall.

[43] Gosden, T.P., Shastri, K.L., Innocenti, P. & Chenoweth, S.F. 2012 The B-matrix harbors significant and sex-specific constraints on the evolution of multicharacter sexual dimorphism. Evolution 66, 2106–2116. (doi:10.1111/j.1558-5646.2012.01579.x.)

[44] Sall, J., Lehman, A. & Creighton, L. 2005 JMP start statistics: a guide to statistics and data analysis Using JMP and JMP IN software. Cary, NC, SAS Institute.

[45] Yang, J., Benyamin, B., McEvoy, B.P., Gordon, S., Henders, A.K., Nyholt, D.R., Madden, P.A., Heath, A.C., Martin, N.G., Montgomery, G.W., et al. 2010 Common SNPs explain a large proportion of the heritability for human height. Nature Genetics 42, 565–569. (doi:10.1038/ng.608.)

[46] Yang, J.A., Lee, S.H., Goddard, M.E. & Visscher, P.M. 2011 GCTA: a tool for genome-wide complex trait analysis. Am. J. Hum. Genet. 88, 76–82. (doi:10.1016/j.ajhg.2010.11.011.)

[47] Patterson, N., Price, A.L. & Reich, D. 2006 Population structure and eigenanalysis. PLoS Gen. 2, 2074–2093. (doi:ARTN e190 10.1371/journal.pgen.0020190).

[48] Falconer, D.S. & Mackay, T.F.C. 1996 Introduction to Quantitative Genetics. 4 ed. Essex, UK, Longman.

[49] Kruijer, W., Boer, M.P., Malosetti, M., Flood, P.J. & Engel, B. 2015 Marker-based estimation of heritability in immortal populations Genetics 200, 385–385.

[50] Kirkpatrick, M. & Meyer, K. 2004 Direct estimation of genetic principal components: simplified analysis of complex phenotypes. Genetics 168, 2295–2306. (doi:10.1534/genetics.104.029181.)

[51] Hine, E. & Blows, M.W. 2006 Determining the effective dimensionality of the genetic variance–covariance matrix. Genetics 173, 1135–1144. (doi:10.1534/genetics.105.054627.)

[52] Zhou, X. & Stephens, M. 2014 Efficient algorithms for multivariate linear mixed models in genome-wide association studies. Nature Methods 11, 407–409. (doi:10.1038/nmeth.2848.)

[53] Lippert, C., Listgarten, J., Liu, Y., Kadie, C.M., Davidson, R.I. & Heckerman, D. 2011 FaST linear mixed models for genome-wide association studies. Nature Methods 8, 833. (doi:10.1038/nmeth.1681 https://www.nature.com/articles/nmeth.1681#supplementary-information.)

[54] Yang, J., Zaitlen, N.A., Goddard, M.E., Visscher, P.M. & Price, A.L. 2014 Advantages and pitfalls in the application of mixed-model association methods. Nature Genetics 46, 100–106. (doi:10.1038/ng.2876.)

[55] Listgarten, J., Lippert, C., Kadie, C.M., Davidson, R.I., Eskin, E. & Heckerman, D. 2012 Improved linear mixed models for genome-wide association studies. Nature Methods 9, 525–526. (doi:10.1038/nmeth.2037.)

[56] Bolstad, G.H., Hansen, T.F., Pelabon, C., Falahati-Anbaran, M., Perez-Barrales, R. & Armbruster, W.S. 2014 Genetic constraints predict evolutionary divergence in Dalechampia blossoms. Philosophical Transactions of the Royal Society of London. Series B - Biological Sciences 369, 20130255. (doi:10.1098/rstb.2013.0255.)

[57] Bai, Z. & Silverstein, J.W. 2010 Spectral Analysis of Large Dimensional Random Matrices. 2 ed. New York, Springer.

[58] Johnstone, I.M. 2006 High dimensional statistical inference and random matrices. arXiv math 0611589.

[59] Benjamini, Y. & Hochberg, Y. 1995 Controlling the false discovery rate: a practical and powerful approach to multiple testing. Journal of the Royal Statistical Society. Series B (Methodological) 57, 289–300.

[60] Yang, J., Weedon, M.N., Purcell, S., Lettre, G., Estrada, K., Willer, C.J., Smith, A.V., Ingelsson, E., O’Connell, J.R., Mangino, M., et al. 2011 Genomic inflation factors under polygenic inheritance. European journal of human genetics 19, 807–812. (doi:10.1038/ejhg.2011.39.)

[61] Reddiex, A.J., Allen, S.L. & Chenoweth, S.F. 2018 A genomic reference panel for Drosophila serrata. G3-Genes Genom Genet 8, 1335–1346.

[62] Skelly, D.A., Magwene, P.M. & Stone, E.A. 2016 Sporadic, Global Linkage Disequilibrium Between Unlinked Segregating Sites. Genetics 202, 427-+. (doi:10.1534/genetics.115.177816.)

[63] Houle, D. & Márquez, E.J. 2015 Linkage disequilibrium and inversion-yyping of the Drosophila melanogaster Genome Reference Panel. G3: Genes|Genomes|Genetics 5, 1695. (doi:10.1534/g3.115.019554.)

[64] Storey, J.D. & Tibshirani, R. 2003 Statistical significance for genomewide studies. Proceedings of the National Academy of Sciences 100, 9440. (doi:10.1073/pnas.1530509100.)

[65] Berner, D. 2012 How much can the orientation of G’s eigenvectors tell us about genetic constraints? Ecology and Evolution 2, 1834–1842. (doi:10.1002/ece3.306.)

[66] Brodie III, E.D. & McGlothlin, J.W. 2007 A cautionary tale of two matrices: the duality of multivariate abstraction. Journal of evolutionary biology 20, 9–14. (doi:10.1111/j.1420-9101.2006.01219.x.)

[67] McGuigan, K., Collet, J.M., McGraw, E.A., Ye, Y.H., Allen, S.L., Chenoweth, S.F. & Blows, M.W. 2014 The nature and extent of mutational pleiotropy in gene expression of male Drosophila serrata. Genetics 196, 911. (doi:10.1534/genetics.114.161232.)

[68] Houle, D. & Fierst, J. 2013 Properties of spontaneous mutational variance and covariance for wing size and shape in Drosophila melanogaster. Evolution 67, 1116–1130. (doi:10.1111/j.1558-5646.2012.01838.x.)

[69] Estes, S., Ajie, B.C., Lynch, M. & Phillips, P.C. 2005 Spontaneous mutational correlations for life-history, morphological and behavioral characters in Caenorhabditis elegans. Genetics 170, 645. (doi:10.1534/genetics.104.040022.)

[70] Chertemps, T., Duportets, L., Labeur, C., Ueda, R., Takahashi, K., Saigo, K. & Wicker-Thomas, C. 2007 A female-biased expressed elongase involved in long-chain hydrocarbon biosynthesis and courtship behavior in <em>Drosophila melanogaster</em>. Proceedings of the National Academy of Sciences 104, 4273–4278. (doi:10.1073/pnas.0608142104.)

[71] Liénard, M.A., Hagström, Å.K., Lassance, J.-M. & Löfstedt, C. 2010 Evolution of multicomponent pheromone signals in small ermine moths involves a single fatty-acyl reductase gene. Proceedings of the National Academy of Sciences 107, 10955–10960. (doi:10.1073/pnas.1000823107.)

[72] Howard, R.W. & Blomquist, G.J. 2005 Ecological, behavioral, and biochemical aspects of insect hydrocarbons. Annual Review of Entomology 50, 371–393.

[73] Delcourt, M., Blows, M.W., Aguirre, J.D. & Rundle, H.D. 2012 Evolutionary optimum for male sexual traits characterized using the multivariate Robertson–Price Identity. Proceedings of the National Academy of Sciences of the United States of America 109, 10414.

[74] Sztepanacz, J.L. & Rundle, H.D. 2012 Reduced genetic variance among high fitness individuals: inferring stabilizing selection on male sexual displays in Drosophila serrata. Evolution 66, 3101–3110. (doi:10.1111/j.1558-5646.2012.01658.x.)

[75] Blows, M.W., Chenoweth, S.F. & Hine, E. 2004 Orientation of the genetic variance-covariance matrix and the fitness surface for multiple male sexually selected traits. American Naturalist 163, 329–340. (doi:10.1086/381941.)

[76] Smith, R.A. & Rausher, M.D. 2008 Selection for character displacement is constrained by the genetic architecture of floral traits in the ivyleaf morning glory. Evolution 62, 2829–2841. (doi:10.1111/j.1558-5646.2008.00494.x.)

[77] Lewis, Z., Wedell, N. & Hunt, J. 2011 Evidence for strong intralocus sexual conflict in the Indian meal moth, Plodia interpunctella. Evolution 65, 2085–2097. (doi:10.1111/j.1558-5646.2011.01267.x.)

[78] Simonsen, A.K. & Stinchcombe, J.R. 2010 Quantifying evolutionary genetic constraints in the ivyleaf morning glory, Ipomoea hederacea. International Journal of Plant Sciences 171, 972–986. (doi:10.1086/656512.)

[79] Teplitsky, C., Tarka, M., Moller, A.P., Nakagawa, S., Balbontin, J., Burke, T.A., Doutrelant, C., Gregoire, A., Hansson, B., Hasselquist, D., et al. 2014 Assessing multivariate constraints to evolution across ten long-term avian studies. PloS one 9, e90444. (doi:10.1371/journal.pone.0090444.)

[80] Walling, C.A., Morrissey, M.B., Foerster, K., Clutton-Brock, T.H., Pemberton, J.M. & Kruuk, L.E.B. 2014 A multivariate analysis of genetic constraints to life history evolution in a wild population of red deer. Genetics 198, 1735–1749. (doi:10.1534/genetics.114.164319.)

[81] Hine, E., McGuigan, K. & Blows, M.W. 2011 Natural selection stops the evolution of male attractiveness. Proc. Natl. Acad. Sci. USA 108, 3659–3664. (doi:10.1073/pnas.1011876108.)

[82] Gosden, T.P., Reddiex, A.J. & Chenoweth, S.F. 2018 Artificial selection reveals sex differences in the genetic basis of sexual attractiveness. Proc. Natl. Acad. Sci. USA 115, 5498–5503. (doi:10.1073/pnas.1720368115.)

[83] Slate, J., Santure, A.W., Feulner, P.G., Brown, E.A., Ball, A.D., Johnston, S.E. & Gratten, J. 2010 Genome mapping in intensively studied wild vertebrate populations. Trends Genet 26, 275–284. (doi:10.1016/j.tig.2010.03.005.)

[84] Ellegren, H. 2014 Genome sequencing and population genomics in non-model organisms. Trends Ecol Evol 29, 51–63. (doi:10.1016/j.tree.2013.09.008.)

